# A compendium of nonredundant short Polymerase III promoters for CRISPR applications

**DOI:** 10.1101/2024.12.25.630128

**Authors:** Michihito Deguchi, Kayla M. Sinclair, Annie Patel, Mckenna Coile, Maria A. Ortega, William P. Bewg, Chung-Jui Tsai

## Abstract

Multiplex genome editing via CRISPR enables simultaneous gene knockout, activation, and repression, but current toolkits often rely on repetitive and lengthy Pol III promoters, limiting construct diversity and genetic stability. Here, we present a diverse collection of minimal U6 and U3 Pol III promoters that support efficient guide RNA (gRNA) expression in dicots. Using CRISPR-Cas9 editing of *mEGFP* in *Nicotiana benthamiana* and *Populus tremula* × *alba*, we demonstrate that minimal promoters can match the performance of their longer counterparts. A systematic evaluation of 14 short promoters revealed that most are functional across species, with exceptions linked to sequence variations in the upstream sequence element (USE) and TATA box. Mutagenesis confirmed the causal role of specific nucleotide changes in promoter activity. Furthermore, we show that regions outside these conserved motifs can be modified to create new-to-nature, functional promoters. This work refines the consensus for functional Pol III promoter elements and expands the toolkit for efficient, stable, and scalable CRISPR editing in dicots, with implications for synthetic promoter design.

---

Multiplex genome editing via CRISPR is a powerful tool for simultaneous knockout, activation, and/or repression of distinct genes. However, current toolkits for multiplex editing lack diversity. Polymerase III (Pol III) promoters are widely used to express guide RNAs (gRNAs). Repeated sequences, including promoters, in multiple expression cassettes complicate construct assembly and have long been a concern for genetic stability and unwanted silencing (Assaad et al., 1993; Peremarti et al., 2010). Expressing gRNAs as a polycistronic array eliminates the need for multiple promoters, but these strategies still involve repeated use of tRNAs, ribozymes, or other RNA-cleaving systems (Xie et al., 2015; Čermák et al., 2017), raising additional concerns about variable gRNA processing. Furthermore, using unnecessarily long promoters may increase the genetic load and introduce uncertainties that impact CRISPR efficiency. Here, we present a diverse collection of short Pol III promoters to support increasingly sophisticated genome editing applications in dicots.

Published plant U6 and U3 promoters range from 79 bp (Nekrasov et al., 2013) to over 700 bp (Ma et al., 2015; Dai et al., 2021) (Supplementary Table S1), with longer promoters being severalfold larger than the transcripts they control. Because the upstream sequence element (USE) and TATA box necessary for Pol III transcription are close to the transcription start site (Waibel and Filipowicz, 1990), we reasoned that minimal Pol III promoters would be effective. We assessed Pol III promoter activity for CRISPR-Cas9 editing of *mEGFP* (monomeric enhanced green fluorescent protein) in stably re-transformed *Nicotiana benthamiana* and poplar (*Populus tremula* × *alba* INRA 717-1B4) reporter lines (Supplementary Text S1). We first tested a deletion series of the experimentally validated *Medicago truncatula MtU6*.*6* promoter (352 bp, Zhou et al., 2015). The 70 bp *MtU6*.*6* promoter was as effective as the 352 bp promoter, based on the loss of mEGFP signal in both root tips and leaf trichomes of *N. benthamiana* (Figure 1A). Amplicon sequencing confirmed that all four promoter lengths had similar efficiencies in *N. benthamiana* and poplar, whereas the promoterless control was ineffective (Figure 1B, Supplementary Table S2).

**Figure 1.**
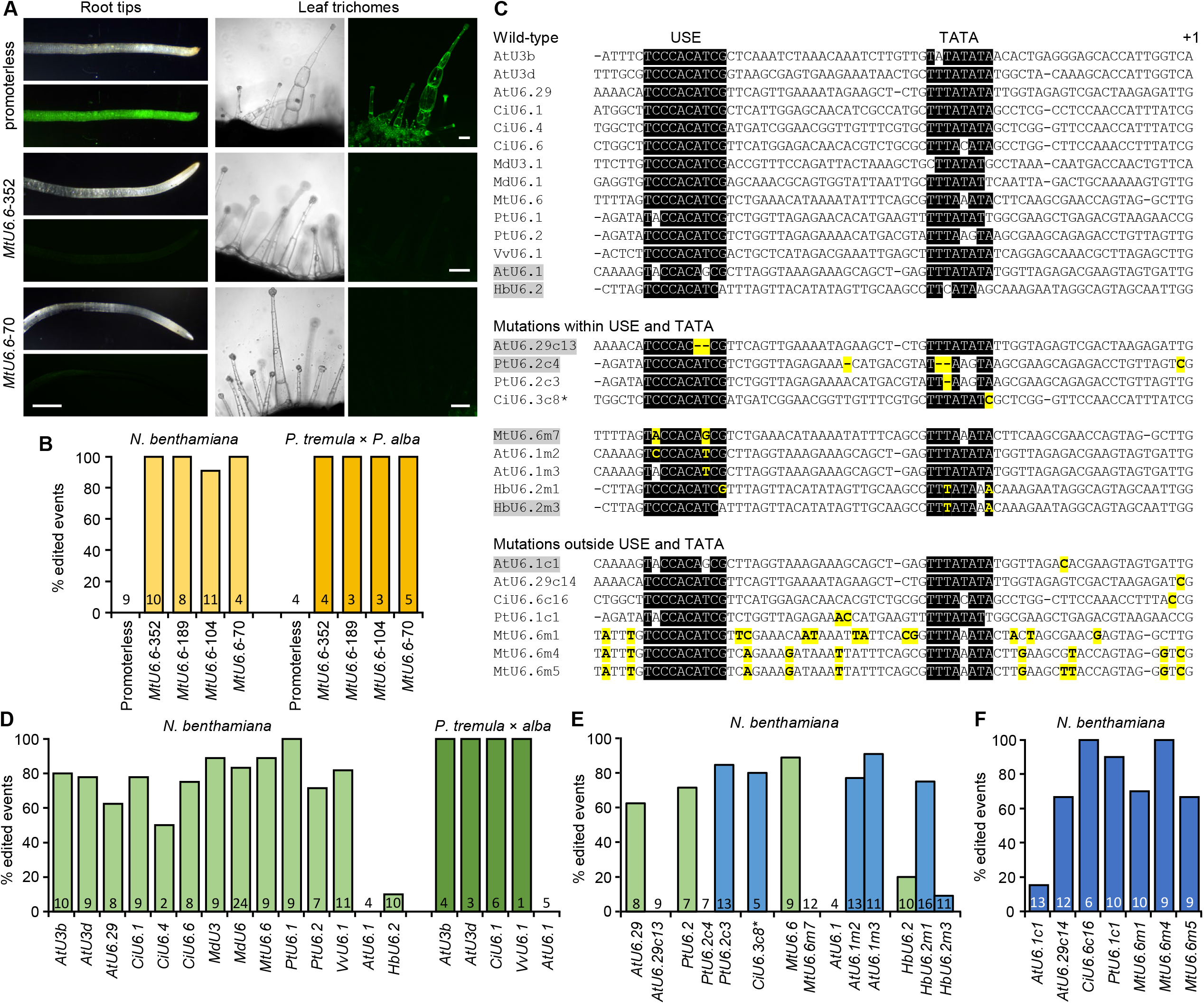
Pol III promoter characterization. A. Images of *N. benthamiana* root tips or leaf trichomes under brightfield or UV. Scale bars: 1 mm (root tips), 100 μm (trichomes). B. Editing efficiencies of the *MtU6*.*6* promoter deletion series. C. Alignments of 70 bp promoters. Conserved USE and TATA sequences are shaded in black, mutations in yellow, and non-functional promoters in grey. D. Editing efficiencies of 70 bp promoters from diverse origins. E. Editing efficiencies of 70 bp promoters with mutations in the USE or TATA (blue bars). Wild-type promoter data (from D, light green bars) are included as reference. F. Editing efficiencies of 70 bp promoters with mutations outside USE and TATA. Transgenic event numbers are indicated at the bottom of the bars. \**CiU6*.*3* wild-type promoter was not tested. At, *Arabidopsis thaliana*; Ci, *Cichorium intybus*; Hb, *Hevea brasiliensis*; Md, *Malus domestica*; Pt, *Populus trichocarpa*; Vv, *Vitis vinifera*.

We next investigated whether the 70 bp length is universally effective for dicot Pol III promoters. We oligo-synthesized 14 short U6/U3 promoters of diverse origin (Figure 1C, Supplementary Table S3) for construct assembly. They were selected with either demonstrated or inconsistent activities (Li et al., 2013; Fan et al., 2015; Ma et al., 2015; Li et al., 2021) or identified from the *Populus trichocarpa* genome with unknown functionality (Supplementary Table S1). All but two (*Arabidopsis thaliana AtU6*.*1* and *Hevea brasiliensis HbU6*.*2*) showed efficient editing in *N. benthamiana* (Figure 1D). The *AtU6*.*1* promoter also failed in poplar, but the other four short promoters tested were functional. These results suggest that the 70 bp length is sufficient for gRNA expression in dicots, with some exceptions.

One of the exceptions, *AtU6*.*1*, was also reported as ineffective in transgenic poplar (Fan et al., 2015; Li et al., 2021), suggesting a genetic basis for the defect. Close examination revealed sequence variations within the USE and/or TATA box of *AtU6*.*1* and *HbU6*.*2* (Figure 1C). To demonstrate causality, we utilized erroneous promoter clones from multiplex Gibson assembly of pooled oligos. We found that 2 nt deletions within the USE (*AtU6*.*29c13*) or TATA box (*PtU6*.*2c4*) abolished editing, whereas a single-base deletion in the TATA box (*PtU6*.*2c3*) was functional (Figure 1E). Mutations outside the conserved elements did not impact activity (Figure 1F). Next, we generated reciprocal mutations between the functional *MtU6*.*6* and the defective *AtU6*.*1* targeting the two variable USE positions. Mutation of the *MtU6*.*6* USE (*MtU6*.*6m7*_*C2A/T8G*_) to resemble *AtU6*.*1* resulted in a complete loss of activity, whereas the reciprocal mutation of *AtU6*.*1* (*AtU6*.*1m2*_*A2C/G8T*_) restored editing (Figure 1E). Correction at the eighth position alone (*AtU6*.*1m3*_*G8T*_) to resemble the USE of *PtU6*.*1* was sufficient to restore activity, indicating the T8G mutation as causal to the *AtU6*.*1* promoter defect. For *HbU6*.*2*, mutations to restore USE (A10G) and TATA box (C3T/G8A) conservation, but not the TATA box alone, rescued promoter activity (Figure 1E). This suggests that the A10G mutation of USE is causal to *HbU6*.*2* inactivity. It is worth noting that some variations in the AT-rich TATA box appeared tolerated, indicative of a less stringent requirement.

The observation that sequences outside the USE and TATA elements are highly variable suggests that non-conserved regions can be exploited to generate novel Pol III promoters. We designed two *MtU6*.*6* promoter variants with 9-13 dispersed substitutions outside the USE and TATA elements (*MtU6*.*6m1* and *MtU6*.*6m4*). A third variant, *MtU6*.*6m5*, was a cloning artifact with an additional substitution compared to *MtU6*.*6m4* (Figure 1C). All three variants retained promoter activity (Figure 1F), supporting the idea that sequences outside USE and TATA can be altered without affecting function.

Finally, we DNA-synthesized a 4-plex design using non-redundant promoters of varying lengths for one-step binary vector cloning to target trichome-regulating *MYB* paralogs in poplar (Figure 2A-B, Supplementary Text S1) (Bewg et al., 2022). Mutations at all target sites were evidenced by the glabrous phenotype and confirmed by amplicon sequencing in transgenic poplar, demonstrating that short promoters are effective in multiplex editing (Figure 2C-D, Supplementary Table S2). This simple and cost-effective approach of synthesizing multi-gRNA cassettes with short, non-redundant U6/U3 promoters for multiplex construct assembly has also been reported elsewhere (Ortega et al., 2023; Nagy et al., 2025).

**Figure 2.**
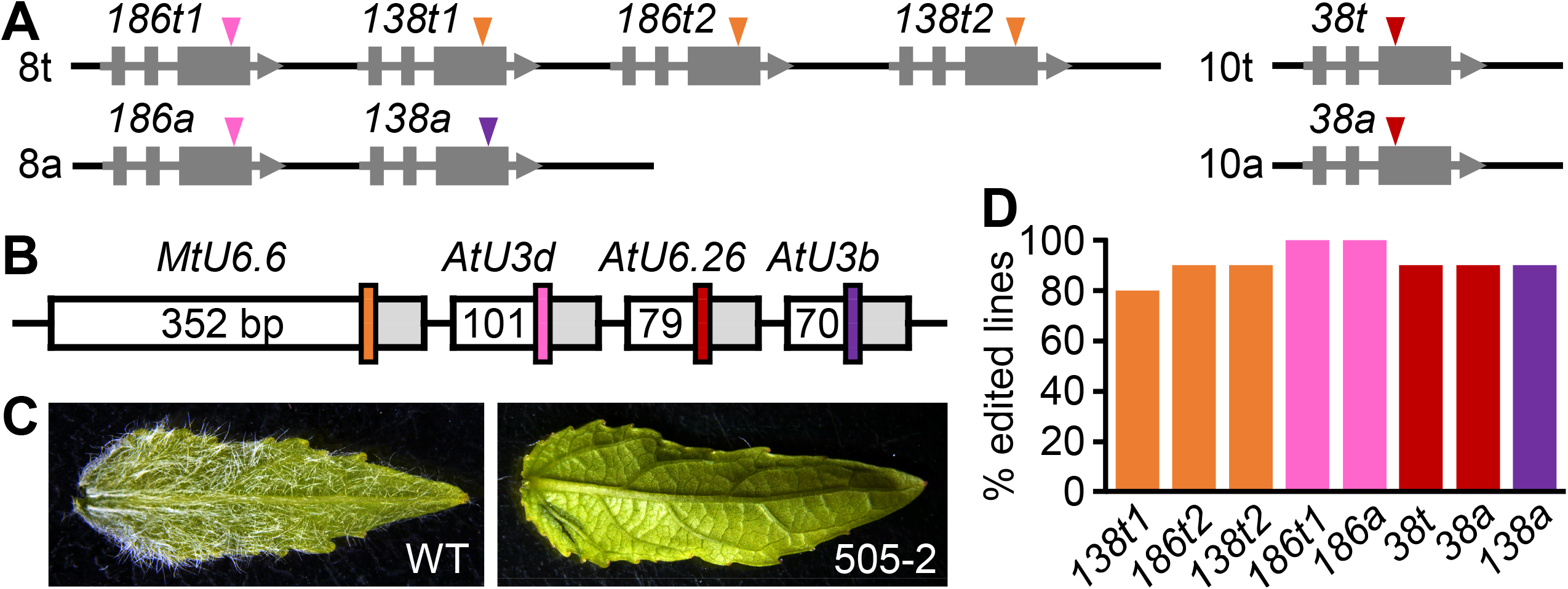
Multiplex editing using nonredundant promoters. A. Schematic representation of the target *MYB186, MYB138*, and *MYB38* alleles on chromosomes 8 and 10 of the *P. tremula* (t) and *P. alba* (a) subgenomes. B. Diagram of the construct design showing four gRNAs driven by distinct promoters of varying lengths (bp). C. Representative leaf images of WT and an edited plant. D. Editing efficiencies at the eight target alleles across 10 transgenic lines, with alleles color-coded by gRNA as in panels A-B.

In summary, this study presents several advances and lessons regarding Pol III promoters. We demonstrate the effectiveness of minimal U6 and U3 promoters from various dicot species. The USE consensus can be refined as TYCCACATCG, whereas the TATA consensus is less strict. Not all wild-type Pol III promoters are functional due to USE variation, as shown for *AtU6*.*1* and *HbU6*.*2*. The updated USE and TATA consensus, along with the mutagenesis data presented, can guide the selection of Pol III promoters with broad functionality. We also exemplified the design of new-to-nature promoters to increase diversity. Given the non-conserved sequence space and combinatorial nucleotide possibilities, the potential for synthetic Pol III promoters is immense. Although this work focused on dicots, similar approaches are applicable to monocot Pol III promoters, which harbor additional monocot-specific *cis* elements upstream of USE (Hao et al., 2020; Nagy et al., 2025). The compendium of 23 experimentally verified dicot U6/U3 promoters should significantly enhance multiplex genome editing capability for CRISPR applications.

## Acknowledgment

We thank Margot Chen for plant transformation assistance, the Georgia Genomics and Bioinformatics Core (RRID:SCR_010994) for Illumina sequencing, and the Biomedical Microscopy Core for Zeiss LSM-880 imaging.

## Author contributions

C.-J.T. conceived the research, C.-J.T and M.D. designed the experiments, M.D., K.S., and A.P. executed the experiments with assistance from M.C., M.A.O. and W.P.B., M.D. and C.-J.T. wrote the paper.

## Supplementary Data

**Supplementary Text S1**. Materials and Methods

**Supplementary Table S1**. Published U6 and U3 promoters from dicot species.

**Supplementary Table S2**. Amplicon sequencing results of transgenic lines.

**Supplementary Table S3**. Oligo and DNA sequences used in this study.

